# Dysmorphic neuron density underlies intrinsic epileptogenicity of the centre of cortical tubers

**DOI:** 10.1101/621607

**Authors:** Sarah EM. Stephenson, Harley G. Owens, Kay L. Richards, Wei Shern Lee, Colleen D’Arcy, Sarah Barton, Simone A. Mandelstam, Wirginia J. Maixner, Duncan MacGregor, Steven Petrou, Paul J. Lockhart, A. Simon Harvey, Richard J. Leventer

## Abstract

Cortical tubers are benign lesions that develop in patients with tuberous sclerosis complex (TSC), often resulting in drug-resistant epilepsy. Surgical resection may be required for seizure control, but the extent of the resection required is unclear. Many centres include resection of perituberal cortex, which may be associated with neurological deficits. Also, patients with tubers in eloquent cortex may be excluded from epilepsy surgery.

Our electrophysiological and MRI studies indicate that the tuber centre is the source of seizures, suggesting that smaller resections may be sufficient for seizure control. Here we report five epilepsy surgeries in four children with TSC and focal motor seizures from solitary epileptogenic tubers in the sensorimotor cortex in whom the resection was limited to the tuber centre, leaving the tuber rim and surrounding perituberal cortex intact. Seizures were eliminated in all cases, and no functional deficits were observed. On routine histopathology we observed an apparent increase in density of dysmorphic neurons at the tuber centre, which we confirmed using unbiased stereology which demonstrated a significantly greater density of dysmorphic neurons within the resected tuber centre (1951 ± 215 cells/mm^3^) compared to the biopsied tuber rim (531 ± 189 cells/mm^3^, n = 4, p = 0.008).

Taken together with our previous electrophysiological and MRI studies implicating the tuber centre as the focus of epileptic activity, and other electrophysiological studies of dysmorphic neurons in focal cortical dysplasia, this study supports the hypothesis that dysmorphic neurons concentrated at the tuber centre are the seizure generators in TSC. Furthermore, our results support limiting resection to the tuber centre, decreasing the risk of neurological deficits when tubers are located within eloquent cortex.

## Introduction

Tuberous sclerosis complex (TSC) is an autosomal dominant, multisystem disorder caused by mutations in the genes encoding hamartin (*TSC1)* or tuberin (*TSC2*). Loss of functional hamartin or tuberin results in failure to form a critical negative regulatory protein complex in the mammalian target of rapamycin (mTOR) pathway (van Slegtenhorst et al., 1998). mTOR overactivation causes dysregulation of neural stem cell proliferation, differentiation, migration and apoptosis, leading to cortical malformation (Aronica and Crino 2014).

TSC is a model for understanding epilepsy due to focal cortical dysplasia (FCD), due to many clinical, histopathological and neuroimaging similarities, particularly with FCD type IIb (Crino 2007). Recent genetic studies of individuals with FCD have identified pathogenic variants in genes encoding multiple components of the mTOR pathway, including *TSC1/2* (Baulac et al., 2015, D’Gama et al., 2015, Leventer et al., 2015, Lim et al., 2015, Scerri et al., 2015, Mirzaa et al., 2016, Sim et al., 2016, D’Gama et al., 2017, Lim et al., 2017). Furthermore, single-cell electrophysiological studies of dysmorphic neurons and balloon cells, the pathognomonic cell lines in FCD type IIb and TSC (Najm et al., 2018), have shown similar electrophysiological properties in both conditions (Cepeda et al., 2005, Cepeda et al., 2010).

Epileptic seizures occur in up to 90% of patients with TSC, often commencing during infancy, having a focal basis, being drug-resistant, and associated with neurodevelopmental impairments (Chu-Shore et al., 2010). Epilepsy surgery is performed in some patients with TSC and drug-resistant epilepsy (DRE), a variety of approaches being employed in seizure localisation and cortical resection, and favourable seizure outcome being reported in the majority (Harvey 2019).

The underlying pathological substrate and cellular mechanisms of epileptogenesis in TSC are debated, being attributed to either cortical tubers, occult dysplasia in perituberal cortex, molecular mechanisms related to the mTORopathy, or pervasive epileptic network disturbances (Wong 2008, Harvey 2019). Our MRI and electrocorticography (ECoG) studies in children with TSC undergoing epilepsy surgery suggested that tubers, not perituberal cortex, are the source of focal seizures in TSC (Mohamed et al., 2012), and that epileptogenicity resides in the centre, not the rim, of epileptogenic tubers (Kannan et al., 2016). When reviewing the histopathology of resected tubers, we repeatedly observed more dysmorphic neurons in the centre compared to the rim of tubers. Additionally, we recognised histopathological and ECoG similarities with bottom-of-sulcus dysplasia (Harvey et al., 2015) and considered performing similar, small, targeted resections in TSC.

Here we provide proof of principle, reporting five surgeries in four children with TSC in whom resection of an epileptogenic tuber in primary motor cortex was limited to the dysplastic tuber centre, eliminating the related seizures and EEG abnormalities and avoiding potential motor deficits. To investigate our hypothesis of an increased density of dysmorphic neurons at the centre of tubers, we performed unbiased stereology in the resected tissue and showed that the density of dysmorphic neurons is significantly greater in the tuber centre than the rim. Our results provide further evidence for epileptogenicity residing in the centre of tubers, further implicate dysmorphic neurons as the generator of seizures in TSC and FCD, and suggest that limited resection may be a practical approach in certain settings.

## Materials and methods

The Royal Children’s Hospital Human Research Ethics Committee approved this study (HREC 29077) and informed consent was obtained for all participants in accordance with the Declaration of Helsinki.

### Clinical details

TSC was diagnosed clinically and confirmed genetically by identification of pathogenic germline variants in *TSC2* in the four children. All had DRE with unilateral tonic or clonic motor seizures, beginning in infancy and occurring at multiple daily frequency at the time of surgery. Three children developed infantile spasms, promptly and effectively treated with vigabatrin; patient 3 was treated with vigabatrin before seizure onset and did not develop infantile spasms. Scalp EEG showed focal interictal epileptiform discharges (IEDs) and ictal rhythms in the contralateral central region. One or more candidate tubers were visible on MRI in the central region.

The children were aged 9 months to 2.9 years at the time of surgery, which was performed between April 2012 and November 2016. Patient 3 underwent separate surgeries for independent right and left-sided focal motor seizures. In all patients, surgery was performed in one stage with intraoperative ECoG, targeting candidate tubers exposed following a stereotactic craniotomy. The precentral and postcentral gyri and adjacent sulci were identified from cross-referencing the cortical surface appearance with their 3D surface-rendered MRI (Figure 1). The candidate tubers, and their centres and sulcal boundaries, were identified in a similar fashion, supplemented by cortical palpation; the concept and boundaries of tuber centres and rims are reported (Kannan et al., 2016). One or two 4-contact depth electrodes (5 mm spaced, Ad-Tech Medical, Wisconsin USA) were used to record from the tuber centres, and either subdural strip (Patients 3a, 3b and 4) or strip and grid (Patients 1 and 2) electrodes (10 mm spaced, Ad-Tech Medical, Wisconsin USA) were used to record from the tuber rims and perituberal cortex. ECoG was recorded from 1-5 candidate tubers in the central and surrounding regions. Runs of rhythmic periodic IEDs or electrographic seizures recorded from the tuber centre indicated their epileptogenicity. EcoG findings and resection details are described in the Results.

**Figure 1.**
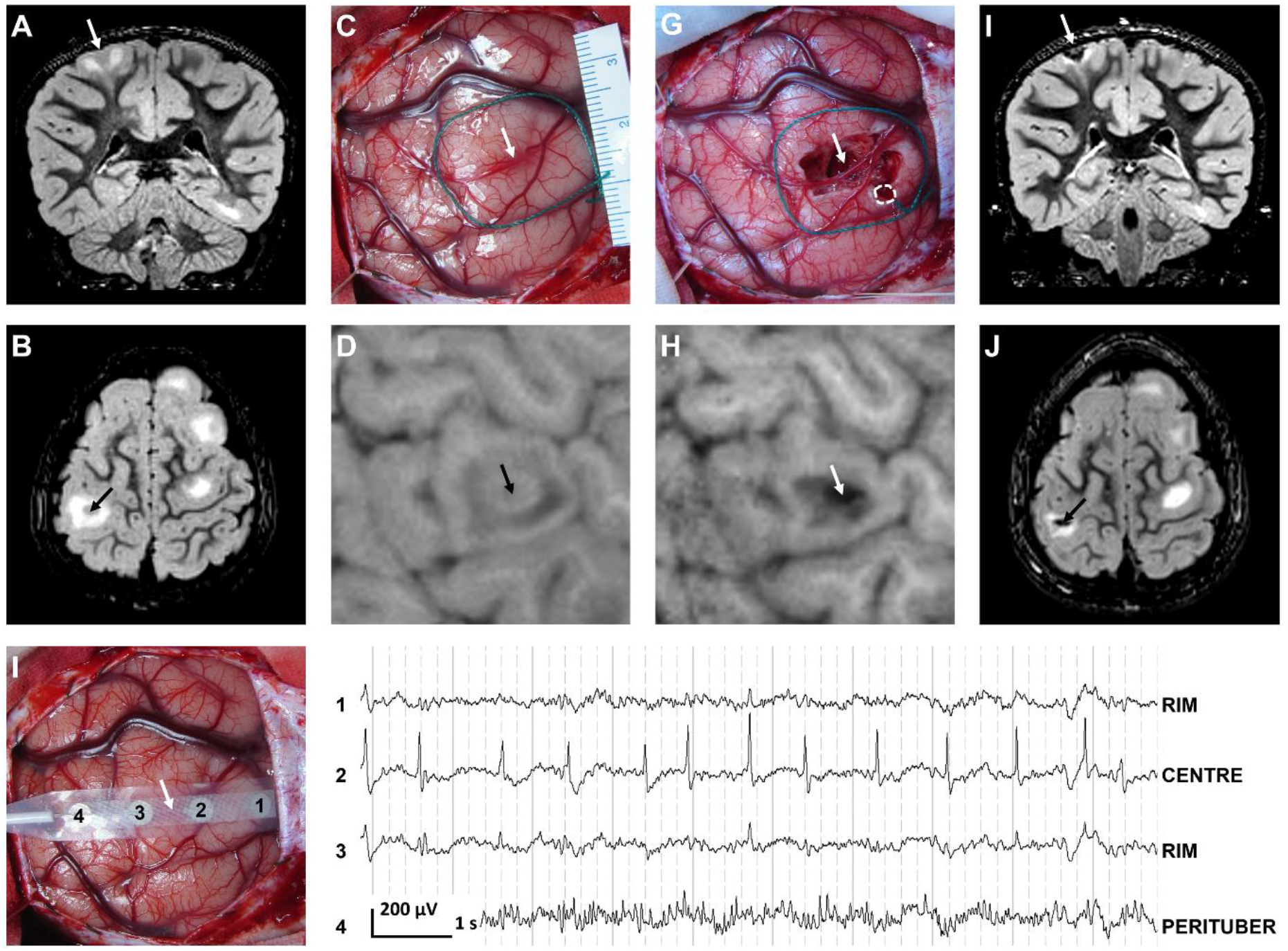
Perioperative MRI scans, photographs and electrocorticography (ECoG) of patient 2, highlighting resection at age 2.8 years of the dysplastic and epileptogenic centre of a left postcentral gyrus tuber causing negative motor seizures of the right arm and epileptic spasms. Preoperative (A and B) and postoperative (I and J) MRI scans in coronal (A and I) and axial (B and J) planes are double inversion recovery sequences acquired at ages 2.8 and 5.8 years, shown in anatomical orientation with the patient’s left side on the left, to correlate with the preoperative (C and D) and postoperative (G and H) photos (C and G) and 3D surface-rendered T1-weighted MRI images (D and H) in which the patient’s left side is on the left and midline is on the right. The preoperative MRI scans (A and B) show thickened cortex with slightly lower signal in the tuber centre, surrounded by high signal in the subcortical white matter of the tuber rim. The preoperative photograph (C) shows the tuber in the postcentral gyrus, identified by correlation of brain surface anatomy with the surface-rendered MRI (D) and palpation, the tuber rim being outlined on the photos by green string and the tuber centre visible as a slight depression. Pre-resection ECoG shown on a referential montage (F), recorded with a 4-contact strip (10mm spacing) straddling the tuber (E), shows continuous, rhythmic spiking at about 1 Hz frequency in the tuber centre contact 2, occasionally propagating to the tuber rim contacts 1 and 3 but not the perituberal cortex contact 4 (F); depth electrode recording was performed separately but was unnecessary, as the tuber centre was shallow and easily recorded with the subdural strip as shown here. The postoperative photograph (G) shows resection of the tuber centre, performed to the grey-white junction at the depth, with a small biopsy of the tuber rim (white dotted circle). The postoperative MRI scans (H, I and J) show a small cavity in the tuber centre surrounded by the high signal of the white matter in the remaining tuber rim. Arrows indicate the tuber centre throughout.

Patient 3 underwent an ineffective resection of a non-epileptogenic left parietal tuber, due to false localisation of her right-sided focal motor seizures, prior to the eventually effective left precentral gyrus tuber-centre resection. Patients 1, 2 and 4 developed independent, non-motor seizures from remote foci that were successfully resected in Patients 1 and 2.

Clinical, EEG, imaging, ECoG and surgical details are summarised for the central surgeries in Table 1 and for all surgeries in Supplementary Table 1.

**Table 1:**
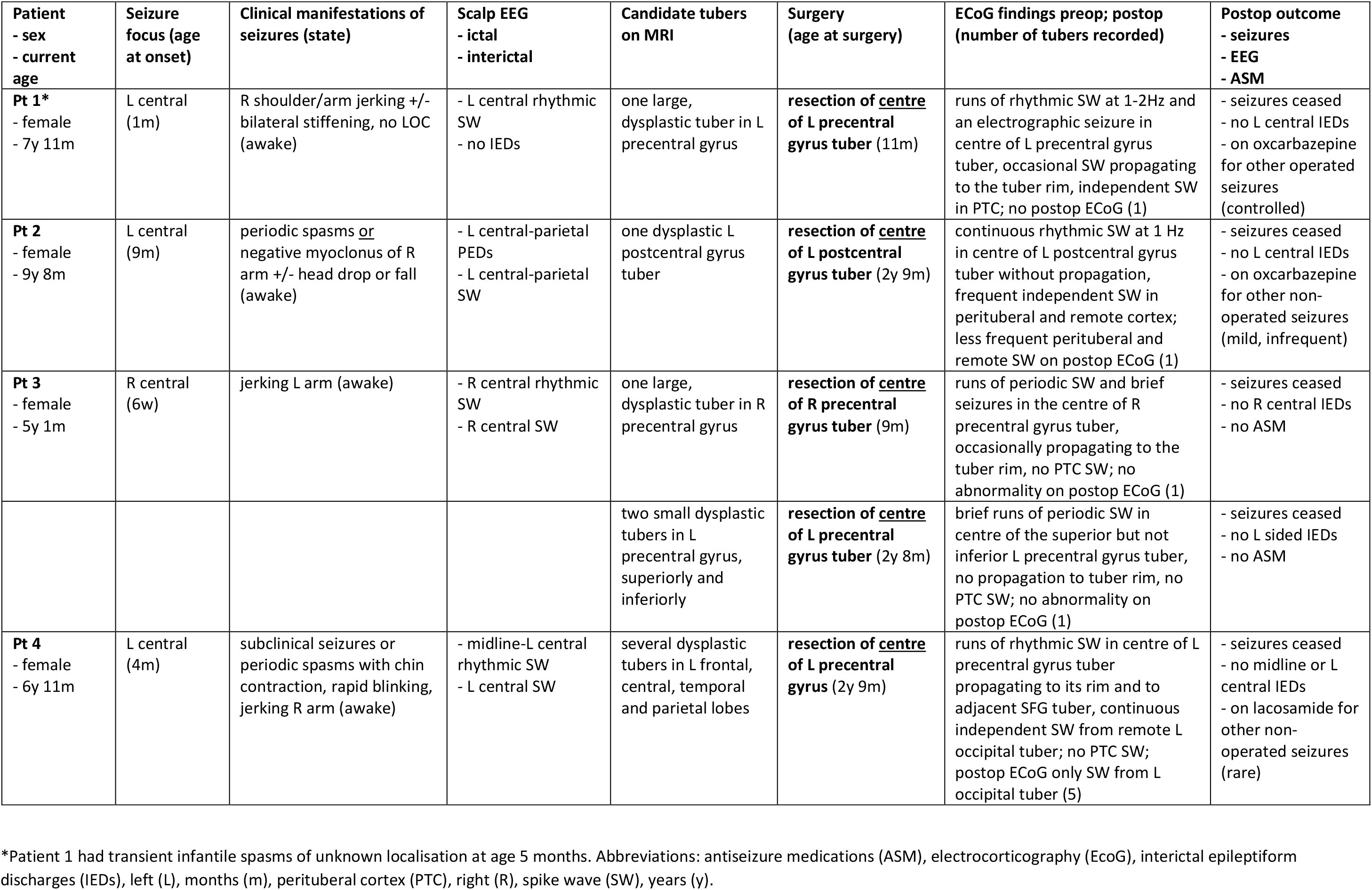
Outline of seizures and tuberectomies performed in patients.

### Tuber stereology details

Tissue samples were collected from the centre (resected tissue) and rim (biopsy) of the operated tuber in patients 2, 3 and 4; no rim biopsy was obtained in patient 1 who was omitted from the stereology study. These were processed for routine histopathological examination. Tissue was placed in formalin, embedded in paraffin, stained with H&E and NeuN, and then reviewed by conventional light microscopy (Supplementary Methods).

Approximately 2 mm of residual paraffin embedded specimen was later cut at 15 µm and 1:10 serial sections were stained with a monoclonal anti-neurofilament antibody (Covance, SMI-311R) to identify dysmorphic neurons (Supplementary Methods) and counterstained with haematoxylin. Stereological analysis of dysmorphic neuron number was performed on blind-coded slides to prevent interpreter bias. Dysmorphic neuron density estimates were obtained using the optical fractionator method (West et al., 1991) via stereology software (StereoInvestigator, Version 11.08.1 [64-Bit]; MicroBrightField (MBF) Bioscience, Williston, Vermont, USA) and a 40x oil immersion lens on a light microscope (DMBL2; Leica, Germany). Dysmorphic neurons were only included in the count if they met the ILAE’s criteria (Blumcke et al., 2011) which required cytoplasmic accumulation of neurofilament proteins, whole cell size measured >20 μm, nuclei measured >15 μm, and abnormally distributed intracellular Nissl substance. For each sample, dysmorphic neurons were counted in every 10th section cut and stained, giving a sampling fraction (ssf) of 1/10. A trace was drawn around the boundaries of the tissue on each section at 5x, within which a counting frame grid was overlaid. The size of each counting frame for the optical dissector probe was 150 μm x 150 μm, giving an area of the counting frame (Aframe) of 22,500 μm^2^. To account for specimen size variability an arbitrary 5% of total surface area per section was sampled. The height of the optical dissector (h) was 10 μm, with an upper guard zone of 2.5 μm. The actual thickness of the mounted sections (t) was measured at five random points within each section analysed.

The estimated total number (N) of dysmorphic neurons in each sample was determined as the sum of cells counted (ΣQ−) on all sections, multiplied by the inverse of the ssf (1/10), the inverse of the area sampling fraction, asf (Aframe/Astep), and the inverse of the thickness sampling fraction, tsf (h/t), according to the formula N = ΣQ− × 1/(ssf × asf × tsf). The coefficient of error (CE) for each stereological estimate of total dysmorphic neuron number was calculated with the Gunderson-Jensen estimator (Gundersen and Jensen 1987) using a smoothness factor of 0 and a = 1/12 (Slomianka and West 2005). The sampling scheme for estimating total dysmorphic neuron number is summarised in Supplementary Table 2. The numerical density (cells/mm^3^; Nv) of dysmorphic neurons was defined as the estimated total number (N) of cells divided by the estimated total volume (Vref) of each sample (given as a default output by StereoInvestigator dictated by the surface area covered by each section trace and the principles of Cavalieri’s basic estimator), according to the formula NV = N/Vref.

### Statistical analysis

For statistical analysis, the paired one-tailed student’s t-test method was employed via Graph Pad Prism (V6, San Diego, USA), where a p value less than 0.05 was considered evidence of statistical significance. Unless otherwise stated, results are expressed as mean ± standard error of the mean (SEM).

## Results

### Tuber centre-only resections

The operated, epileptogenic tubers were in the precentral (Patients 1, 3a, 3b, 4) or postcentral gyrus (Individual 2), abutting the central sulcus. Intraoperative ECoG showed runs (Patients 1, 3a, 3b, 4) or continuous (Patient 2) rhythmic spike-wave activity and electrographic seizures (Patients 1, 3a) in the tuber centre, recorded by depth electrodes. These were confined to the tuber centre (Patients 2, 3b,) or propagated to the tuber rim (Patients 1, 3a, 4). ECoG from the surrounding perituberal cortex showed independent spike-wave activity in Patients 1 and 2. The grey matter of the tuber centre was resected in a subpial fashion using an ultrasonic aspirator, to the grey-white junction. This left a cavity of variable depth in the centre of the tuber, with the cortex of the tuber rim surrounding. The abnormal white matter underlying the cortex of the tuber was not resected. Post-resection ECoG from the tuber rim and perituberal cortex, using subdural strip electrodes, showed no epileptiform activity; independent spiking persisted in the perituberal cortex of Patient 2 (Figure 1, Supplementary Figure 1).

No patient had postoperative weakness, even transiently, and none has a motor or sensory deficit on examination at follow-up. The focal motor seizures ceased immediately following the five surgeries in the four patients, with no recurrence during the 2.4-7.0 (mean 5.0) years of close clinical monitoring. Routine postoperative scalp EEG recordings three months following surgery, and subsequent video-EEG monitoring for independent seizures, did not reveal IEDs or electrographic seizures in the operated central region, except for patient 4 who developed ipsilateral centrotemporal spikes without Rolandic epilepsy (Table 1).

Antiseizure medications (ASM) were ceased following these surgeries in Patients 3 and 4. Patient 3 remains off ASM. Patient 4 was commenced on lacosamide three years later for infrequent, non-motor seizures of remote origin. Patients 1 and 2 had medications reduced following this and subsequent surgeries, to oxcarbazepine monotherapy. No patient has regular, ongoing seizures. Patients 2 and 4 have mild learning and behavioural difficulties, but all attend mainstream schooling. Patient 1 has an intellectually disability and autism spectrum disorder and attends a special developmental school. Patient 3 is developmentally normal.

### Histopathology and dysmorphic neuron density

Histopathological examination of tuber specimens showed similar features in each. Cortical neuronal layering was distorted by dysmorphic and enlarged neurons and balloon cells, resulting in blurring of the cortical-white matter junction. On routine histopathological examination using H&E staining (Figure 2) and NeuN immunohistochemistry (data not shown), the density of dysmorphic neurons appeared greater in the tuber centre of cortical specimens relative to the tuber rim biopsies, the latter showing a predominance of balloon cells and fewer dysmorphic neurons.

**Figure 2.**
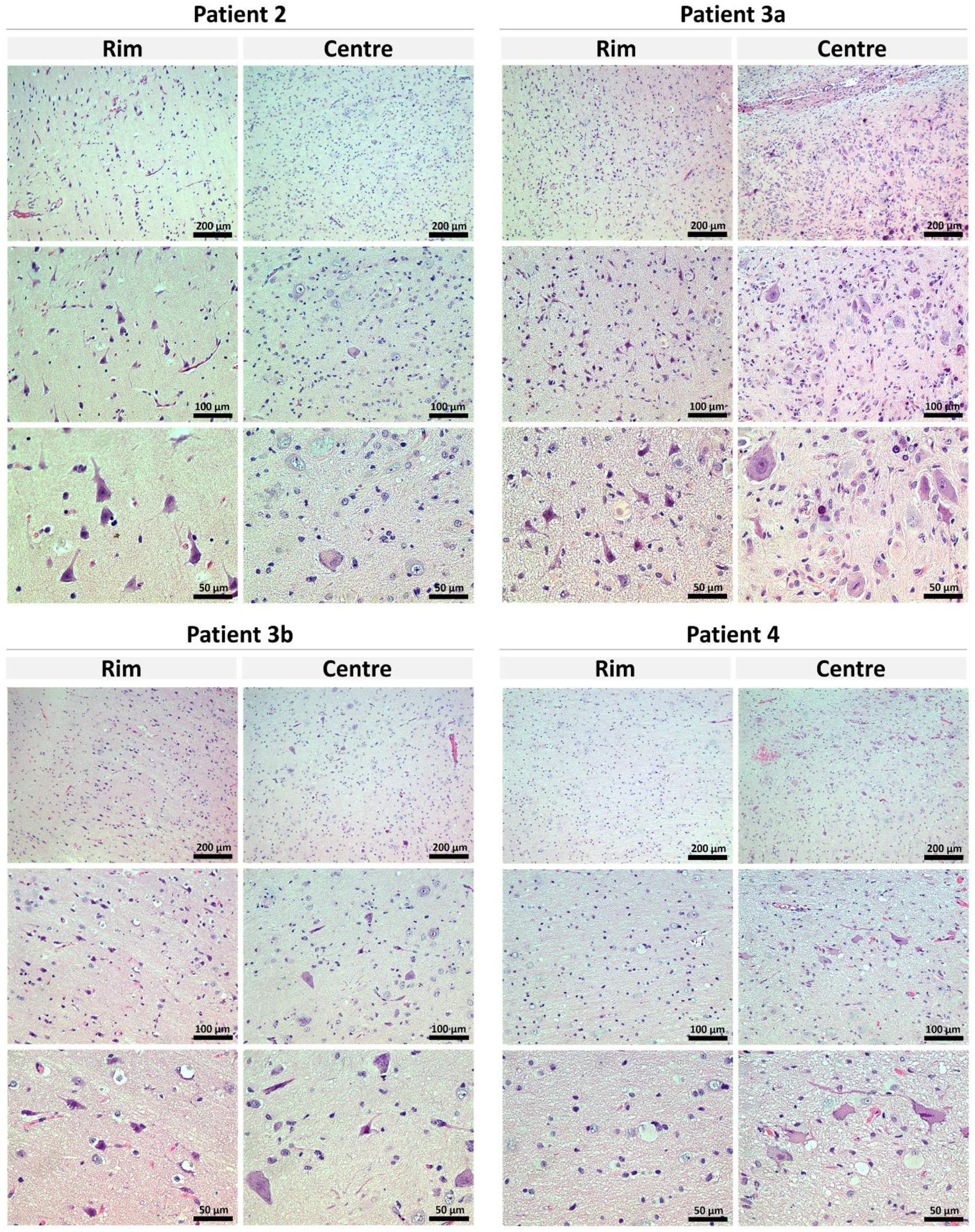
Qualitative differences in dysmorphic neurons numbers identified in specimens from the centre of cortical tubers compared to the rim. Haemotoxylin and Eosin staining from paired biopsies taken from the rim and the centre of cortical tubers demonstrate spatial differences in the laminal architecture and cellular constituents within specimens. Specimens resected from the tuber centre also largely comprised of grey matter but demonstrated marked dyslamination with irregular clusters of large, cytomegalic and dysmorphic neurons and balloon cells. Conversely, biopsy specimens from the tuber rim largely comprised grey matter in which cortical lamination appeared relatively normal, cytomegalic or dysmorphic neurons were rarely identified, and there was a predominance of balloon cells. Scale bars are indicated.

Stereological measurement demonstrated a greater density of dysmorphic neurons within the tuber centre (1951 ± 215 cells/mm^3^, mean ± SEM, n=4) compared to the rim (531 ± 189 cells/mm^3^, mean ± SEM, n=4) (p=0.008) (Fig. 3). The significantly higher density of dysmorphic neurons at the centre compared to the rim was present in each individual when analysed independently, with a mean difference of 1420 cells/mm^3^ (95% confidence interval 490 to 2351).

**Figure 3.**
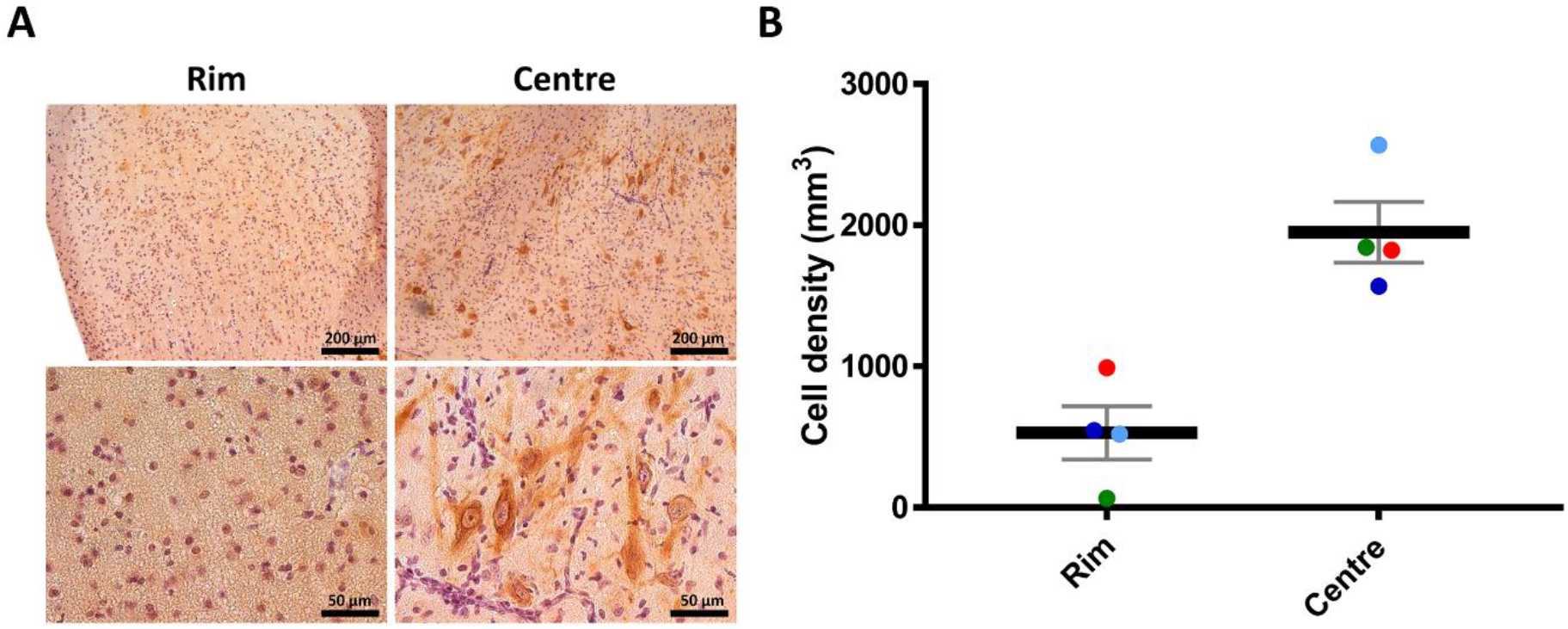
The density of dysmorphic neurons is greater in the centre of the tuber. (A) Representative images of anti-neurofilament protein immunohistochemistry staining the dysmorphic neurons in tuber rim and tuber centre demonstrating a typically observed cluster of dysmorphic neurons in the centre biopsy that were absent from the rim. Scale bars are indicated. (B) Unbiased quantitative stereology determined the density of dysmorphic neurons in the tuber centre was greater than the tuber rim. Patient 2 – green; Patient 3a – light blue; Patient 3b – dark blue; patient 4 – red. Cell densities (expressed as cell number/mm3) on the y-axis are compared between tuber rim and centre on the x-axis for each paired sample. Error bars represent standard error of the mean.

## Discussion

The approach to epilepsy surgery in TSC varies significantly between hospitals. Presurgical workup is complex and involves detailed clinical, imaging and electrophysiological studies to identify the epileptogenic tubers and best approach for their removal. Surgery is considered early in children with DRE to minimise the detrimental effects of seizures, EEG discharges and ASM on the developing brain. However, some children are not considered candidates for epilepsy surgery due to their epileptogenic tubers being in eloquent cortex, seizures being poorly localised, or tuber load being high.

These four children with tuber centre resections were selected from our surgical series because of the localisation of their seizures. The certainty of seizure remission was clearer clinically than in many children with TSC, given their epileptogenic tubers were located in sensorimotor cortex. Additionally, this was the patient group in which we first undertook this minimalistic approach, to minimise potential injury to surrounding motor cortex and corticospinal tracts. Notably, only a single tuber was found to be responsible and resected, and follow-up was lengthy after surgery. We have employed this technique in several other TSC patients not reported here, those being children with poorly localised usually non-motor seizures and large tuber loads in whom the centres of several tubers were resected, limiting the amount of tissue resected.

Electrophysiological studies of the seizure generating zone in TSC have shown variable results, with some suggesting seizure onset within perituberal cortex (Madhavan et al., 2007, Major et al., 2009, Marcotte et al., 2012, Chalifoux et al., 2013, Ruppe et al., 2014, Sosunov et al., 2015) and others within tubers (Ma et al., 2012, Mohamed et al., 2012, Kannan et al., 2016, Despouy et al., 2019). Many studies lacked detailed methods to determine the spatial and temporal characteristics of seizure onset and spread within, around and between tubers (see review in Harvey 2019). The view that seizures originate in perituberal cortex or widely distributed networks potentially leads to large resections or exclusion of patients from surgery, possibly inappropriately. Our intracranial EEG monitoring studies in children with TSC showed that focal seizures and IEDs arise from the centre of epileptogenic tubers and propagate to the tuber rim, perituberal cortex and other tubers (Mohamed et al., 2012, Kannan et al., 2016). A recent single patient study using new hybrid electrodes with tetrodes demonstrated the tuber is critically involved in seizure onset, with a gradient of epileptogenicity from the tuber centre to the perituberal cortex (Despouy et al., 2019). These electrophysiological findings correlate with the features of dysplasia on MRI and histopathology being more marked in the tuber centres (Kannan et al., 2016).

There are limited descriptions of the microstructure and histopathology of cortical tubers, and few of dysmorphic neuron distribution (Muhlebner et al., 2016a, Muhlebner et al., 2016b). On review of routine histopathology in our wider TSC surgical series, we repeatedly observed abundant dysmorphic neurons in tuber centre specimens and hypothesised that this might be the cause of the increased epileptogenicity at the tuber centre that we detected with our electrophysiology studies. Therefore, we performed objective, quantitative analysis using stereological methods to assess the density of dysmorphic neurons in cortical tubers and demonstrated that the tuber centre contains 3.7 fold greater density of dysmorphic neurons than the rim. We suggest that the efficacy of limited tuber centre resections in these patients is due to removal of a nidus of dysmorphic, and intrinsically epileptic, neurons.

It is suggested that dysmorphic neurons are the pathogenic substrate driving epileptogenesis in FCD, being the source of ictal discharges (Chassoux et al., 2000, Battaglia et al., 2013). Compared to normal pyramidal neurons, dysmorphic neurons possess atypical passive membrane properties. Due to their increased cell membrane area, dysmorphic neurons exhibit a larger cell capacitance, a longer time constant and a lower input resistance (Cepeda et al., 2003). Additionally, dysmorphic neurons are predisposed to hyperexcitability via larger amplitudes of macroscopic Ca2+ currents and Ca2+ influx. Studies demonstrating high expression of ionotropic and metabotropic glutamate receptor subtypes and/or subunits in these cells also provide evidence of dysfunctional molecular pathways contributing to hyperexcitability (White et al., 2001, Wang et al., 2007, Boer et al., 2008). Conversely, balloon cells have a high input resistance, do not exhibit voltage-gated Na+ or Ca2+ currents, and do not generate spontaneous synaptic currents (Cepeda et al., 2003). Further to this, studies demonstrating an insensitivity of balloon cells to excitatory amino acids suggest a low epileptogenic potential (Cepeda et al., 2012), and studies demonstrating increased clearance of glutamate in areas with a high density of balloon cells suggest a protective or antiseizure propagation function (Gonzalez-Martinez et al., 2011). This is of potential relevance to our surgical approach, in which a tuber rim rich in balloon cell remains between the tuber centre resection cavity and normal perituberal cortex.

Our study provides strong evidence for hyperexcitable, dysmorphic neurons concentrated in the centre of cortical tubers being the pathological substrate of DRE in TSC. Our recent report of second-hit *DEPDC5* somatic mutation in a child with FCD being limited to dysmorphic neurons in the seizure focus (Lee et al., 2019) suggests that localised, cell-specific mechanisms may underlie epilepsy in other forms of dysplasia. In bottom-of-sulcus dysplasia, dysmorphic neurons are concentrated at the depth of the dysplastic sulcus where MRI and ECoG abnormalities are greatest, and seizures are similarly controlled with small, targeted resections (Harvey et al., 2015, Jackson et al., 2017).

Limited, tuber-centre resections in children with TSC might be considered in settings other than epileptogenic tubers in eloquent cortex. Given the multiplicity of seizure foci, the large tuber load, the involvement of homologous brain regions and the difficulties with seizure localisation in many children with TSC, multiple tubers could be safely operated using this technique, without the need for resecting large volumes of cerebral tissue. Additionally, laser interstitial thermal therapy is being employed in TSC (Tovar-Spinoza et al., 2018) but could be directed at tuber centres, rather than ablating whole tubers, reducing the risk of thermal injury to normal perituberal cortex.

## Acknowledgements

The authors thank the families for agreeing to be part of this research study. We thank Greta Gillies and Kate Pope for their contributions.

## Funding Information

This work was supported by the Australian Government National Health and Medical Research Council GNT1128933, the Murdoch Children’s Research Institute, the Campbell Edwards Trust and the Brain Foundation. Additional funding was provided by the Independent Research Institute Infrastructure Support Scheme and the Victorian State Government Operational Infrastructure Program. RJL is supported by the Melbourne Children’s Clinician Scientist Fellowship scheme.

