## Supplementary for "Dysmorphic neuron density underlies intrinsic epileptogenicity of the centre of cortical tubers"

**Supplementary Methods**

**Tissue fixation, embedding and sectioning**

All the initial tissue sample processing and microtome cutting steps were performed by the Anatomical Pathology Department at RCH. After resection, tissue was briefly fixed by incubation in 4% (w/v) paraformaldehyde in phosphate buffer for a minimum of 60 min at 4°C, before being transferred to ice-cold 70% (v/v) ethanol. Tissue samples were then dehydrated and perfused with paraffin wax using a Shandon Citadel 1000 Tissue Processor with the following cycle program: formalin for 100 minutes, ethanol for 30 minutes, ethanol for 30 minutes, ethanol for 30 minutes, ethanol for 60 minutes, ethanol for 90 minutes, ethanol for 80 minutes, xylene for 45 minutes, xylene for 75 minutes, xylene for 80 minutes and paraffin wax three times for 60 minutes. Samples were then transferred to a sample mould, overlaid with wax and solidified on a cold-plate at -5°C. Sections were cut at 3-8 μm for routine histology or 15 μm for stereology using a Leica RM2125RT microtome, and floated in a 37°C water bath for 1-5 minutes to remove folds in the wax. The sections were then transferred onto Superfrost+ positively-charged slides (Menzel Gläser) and dried at 37°C for 60 minutes. Slides were stored at room temperature until needed.

**Hematoxylin and eosin stain**

Routine histology slides were stained using the Leica Autostainer and Coverslipper as follows: xylene for three minutes, xylene for three minutes, twice in absolute ethanol for 1 minute, 95% ethanol for 1 minute, water for one minute, haematoxylin for 3 minutes, water for one minute, Acid Alcohol (70% v/v ethanol, 0.25% v/v hydrochloric acid) for 10 seconds, water for one minute, Scotts Tap water (420 mM Sodium Hydrogen Carbonate, 166 mM Magnesium Sulphate) for one minute, water for one minute, absolute ethanol for one minute, eosin for two minutes, twice in absolute ethanol for one minute, xylene for one minute.

**Immunohistochemistry**

Slides were immersed in histolene twice for 3 minutes and histolene:ethanol (1:1) for three minutes, followed by rehydration in 100% ethanol for three minutes, 95% ethanol for three minutes, 70% ethanol for three minutes, 50% ethanol for three minutes and running cold tap water until well rinsed. Specimens were immersed for 1 hour in a blocking solution containing 10% (v/v) normal goat serum (Sigma-Aldrich, G9023) and 0.3% (v/v) Triton X-100 (Sigma-Aldrich, T8787) made up in 0.1 M phosphate buffer (PB; 75mM Na2HPO4 [Merck], 25 mM NaH2PO4 [Merck]; pH 7.4) + 0.05% (v/v) Pro-Clin (Sigma-Aldrich). The blocking solution was removed and the slides were incubated with mouse anti-neurofilament monoclonal antibody (1:1500, Covance, SMI-311R) or 1:100 anti-NeuN (Millipore, MAB377) in 50% blocking solution, 50% 0.1 M PB) overnight at 4°C. The slides were washed three times for 10 minutes in 0.1 M PB and subsequently incubated with biotinylated goat anti-mouse secondary antibody (1:300 Vector Labs, BA-9200 in 50% blocking solution, 50% 0.1 M PB) for 1 hour. The slides were then washed three times for 10 minutes in 0.1 M PB, incubated with Avidin-Biotin Complex (ABC; prepared as per manufacturer's instructions; Vector Labs) for 30 minutes and washed again three times for 10 minutes in 0.1 M PB. Slides were immersed in 3% (v/v) hydrogen peroxide solution (Sanofi) for 5 minutes to quench endogenous peroxidases, followed by three 10 minute washes in water. Diaminobenzidine (DAB; prepared as per manufacturer's instructions,

Vector Labs) was applied to the slides to develop a visible brown precipitate label for dysmorphic neurons, with the staining intensity monitored via microscope. When appropriate staining was observed, the slides were washed three times for 10 minutes in 0.1 M PB, counterstained in haematoxylin (Amber Scientific) for 10 seconds, rinsed in water, dipped in acid ethanol (70% ethanol with 0.25% 12M HCl) ten times, rinsed again, and then dehydrated in via an histolene and ethanol series carried out in reverse order of the above rehydration procedure. Once dehydrated, slides were cover-slipped with MicroMount (Leica).

### Supplementary Table 1: Outline of all seizures and all tuberectomies performed in patients.

| **Patient<br>- sex<br>- current age** | **Seizure focus<br>(age at onset)** | **Clinical manifestations of seizures (state)** | **Scalp EEG<br>- ictal<br>- interictal** | **Candidate tubers on MRI** | **Surgery<br>(age at surgery)** | **ECoG findings preop; postop<br>(number of tubers recorded)** | **Postop outcome<br>- seizures<br>- EEG<br>- ASM** |
|---|---|---|---|---|---|---|---|
| **Pt 1***<br>- female<br>- 7y 11m | L central<br>(1m) | R shoulder/arm jerking +/- bilateral stiffening, no LOC (awake) | - L central rhythmic SW<br>- no IEDs | one large, dysplastic tuber in L precentral gyrus | **resection of centre of L precentral gyrus tuber** (11m) | runs of rhythmic SW at 1-2Hz and an electrographic seizure in centre of L precentral gyrus tuber, occasional SW propagating to the tuber rim, independent SW in perituberal and remote L temporal cortex; no postop ECoG (1) | - seizures ceased<br>- no L central IEDs<br>- reduced ASM |
| | *L temporal<br>(12m)* | *staring, fearful, R arm dystonia, pupils dilate, LOC, oral automatisms (asleep)* | *- L temporal PEDs<br>- L temporal SW* | *two large, dysplastic tubers in L lateral and L basal temporal lobe* | *resection of L lateral temporal and L basal temporal tubers (1y 5m)* | *frequent SW over rim and centre of L lateral temporal tuber and PTC (2)* | *- seizures ceased<br>- no L temporal IEDs<br>- reduced ASM* |
| | *L parietal-occipital<br>(3y 9m)* | *axial automatisms, bilateral stiffening, oral automatisms, LOC, eye deviation to L (awake or asleep)* | *- diffuse L RDA<br>- L posterior quadrant slowing, SW and PFA* | *several small tubers in L parietal, posterior temporal and occipital lobes* | *resection of L parietal and L posterior temporal-occipital tubers (7y 2m)* | *intermittent SW in PTC surrounding the parietal and posterior temporal tubers (6)* | *- seizures ceased<br>- persistent L posterior temporal SW, slowing and PFA<br>remains on oxcarbazepine* |
| **Pt 2**<br>- female<br>- 9y 8m | L central<br>(9m) | periodic spasms or negative myoclonus of R arm +/- head drop or fall (awake) | - L central-parietal PEDs<br>- L central-parietal SW | one dysplastic L postcentral gyrus tuber | **resection of centre of L postcentral gyrus tuber** (2y 9m) | continuous rhythmic SW at 1 Hz in centre of L postcentral gyrus tuber without propagation, frequent independent SW in perituberal and remote cortex; less frequent perituberal and remote SW on postop ECoG (1) | - seizures ceased<br>- no L central IEDs<br>- reduced ASM |
| | *L occipital<br>(1y 9m)* | *subclinical or arousal, blinking, vocalisation, wriggling or prolonged R version, vomiting, convulsing (asleep)* | *- L occipital rhythmic SW then slowing<br>- L occipital SW* | *several tubers in L occipital, posterior temporal and parietal lobes* | *resection of L cuneus tuber (5y)* | *continuous SW in centre and rim of L cuneus tuber only, independent SW and fast activity in PTC; persistent SW and fast activity in cuneus (3)* | *- seizures ceased<br>- no L occipital IEDs or ictal rhythms<br>- reduced ASM* |
| | *R temporal<br>(5y)* | *oral/epigastric discomfort, scared, gagging, vomiting, eye deviation to L, no LOC (awake or asleep)* | *- R temporal RDA then rhythmic SW<br>- R temporal SW* | *two dysplastic tubers in basal R temporal lobe* | *resection of two R temporal tubers (6y 6m)* | *Intermittent SW in centre and rim of both R temporal tubers* | - seizures ceased<br>*- no R temporal IEDs<br>- reduced ASM* |
| | *R frontal<br>(1y 8m)* | *fall, version to L, jerking L arm, LOC, postictal L arm weakness (awake)* | *- R frontal RDA then PEDs<br>- R frontal SW* | *several tubers in R superior frontal gyrus* | *not operated* | *N/A* | *continued seizures on oxcarbazepine* |
| **Pt 3**<br>- female<br>- 5y 1m | R central<br>(6w) | jerking L arm (awake) | - R central rhythmic SW<br>- R central SW | one large, dysplastic tuber in R precentral gyrus | **resection of centre of R precentral gyrus tuber** (9m) | runs of periodic SW and brief seizures in the centre of R precentral gyrus tuber, occasionally propagating to the | - seizures ceased<br>- no R central IEDs<br>- reduced ASM |

| | | | | | | | |
|---|---|---|---|---|---|---|---|
| | | | | | | tuber rim, no PTC SW; no abnormality on postop ECoG (1) | |
| | L central<br>(1y 2m) | R hand tonic, giggling, R face weakness (awake) | - L central-parietal RDA<br>- L post temporal-occipital PEDs | *dysplastic tuber in L parietal convexity and L basal temporal lobe* | *resection of L superior parietal lobule tuber* | *intermittent SW over and around superior parietal tuber (2)* | *- continued seizures and L posterior quadrant IEDs led to subsequent surgery* |
| | | | | two small dysplastic tubers in L precentral gyrus, superiorly and inferiorly | **resection of <u>centres</u> of L precentral gyrus tuber** (2y 8m) | brief runs of periodic SW in centre of the superior but not inferior L precentral gyrus tuber, no propagation to tuber rim, no PTC SW; no abnormality on postop ECoG (2) | - seizures ceased<br>- no L sided IEDs<br>- ceased ASM |
| **Pt 4**<br>- female<br>- 6y 11m | L central<br>(4m) | subclinical seizures or periodic spasms with chin contraction, rapid blinking, jerking R arm (awake) | - midline-L central rhythmic SW<br>- L central SW | several dysplastic tubers in L frontal, central, temporal and parietal lobes | **resection of <u>centre</u> of L precentral gyrus** (2y 9m) | runs of rhythmic SW in centre of L precentral gyrus tuber propagating to its rim and to adjacent SFG tuber, continuous independent SW from remote L occipital tuber, no PTC SW; postop ECoG only SW from L occipital tuber (5) | - seizures ceased<br>- no midline or L central IEDs<br>- ceased ASM |
| | *L occipital<br>(14m)* | *subclinical* | *- L occipital RDA and rhythmic SW<br>- L posterior temporal-occipital SW* | *several dysplastic and cystic tubers in L occipital and posterior temporal lobes* | *not operated, spontaneously resolved* | *N/A* | *- no seizures<br>- no L occipital IEDs or seizures* |
| | *L frontal<br>(3y)* | *LOC, hand tremor, odd smile and mouth movements, grunting, dilated pupils, cluster or prolonged (awake)* | *- L frontal PEDs<br>- L centrotemporal spikes* | *several dysplastic tubers in L frontal lobe* | *not operated* | *N/A* | *- rare seizures<br>- commenced lacosamide* |

*Patient 1 had transient infantile spasms of unknown localisation at age 5 months. Details in italics refer to seizures and tubers not reported in this study. Abbreviations: antiseizure medications (ASM), electrocorticography (EcoG), interictal epileptiform discharges (IEDs), left (L), months (m), perituberal cortex (PTC), right (R), spike wave (SW), years (y).

**Supplementary Table 2: Sampling scheme for estimating total dysmorphic neuron number**

| | Centre mean (SEM) | Rim mean (SEM) |
|---|---|---|
| Measured Volume ($mm^3$) | 18 (3) | 5 (1) |
| Mean number of neurons counted in each sample | 118 (13) | 11 (6) |
| Number of Sampling Sites | 307 (51) | 88 (18) |
| Mounted Thickness | 14.4 (0.2) | 14.5 (0.1) |
| Estimated Population | 33775 (3777) | 3376 (1688) |
| Mean coefficient of error (CE) | 0.09 (0.005) | 0.49 (0.18) |
| Mean *tsf*: thickness sampling fraction | 0.70 (0.007) | 0.69 (0.007) |
| *h* (µm): height of optical disector | 10 | |
| Astep ($\mu m^2$) range: area associated with step | 449,999 | |
| *ssf*: section sampling fraction | 0.1 | |
| Aframe ($\mu m^2$): area of counting frame | 22,500 | |

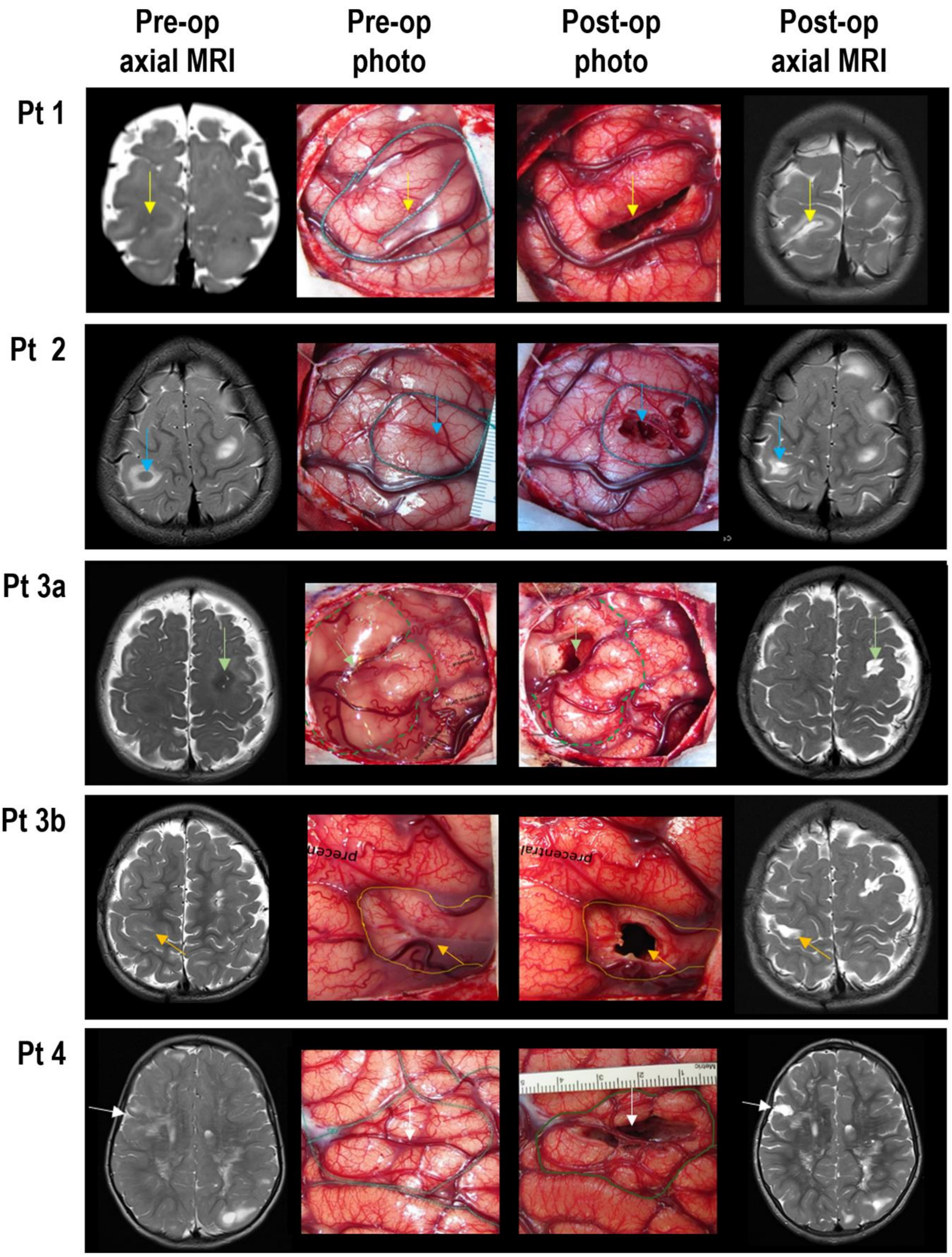


**Supplementary Figure 1: Preoperative and postoperative axial T2-weighted MRI scans and photography.** Anatomical not radiological view i.e. anterior is top and left is left, and operative photos (same orientation as MRI) of the 5 tuber-centre resections in 4 patients. Tuber rims are outlined and tuber centres are arrowed.

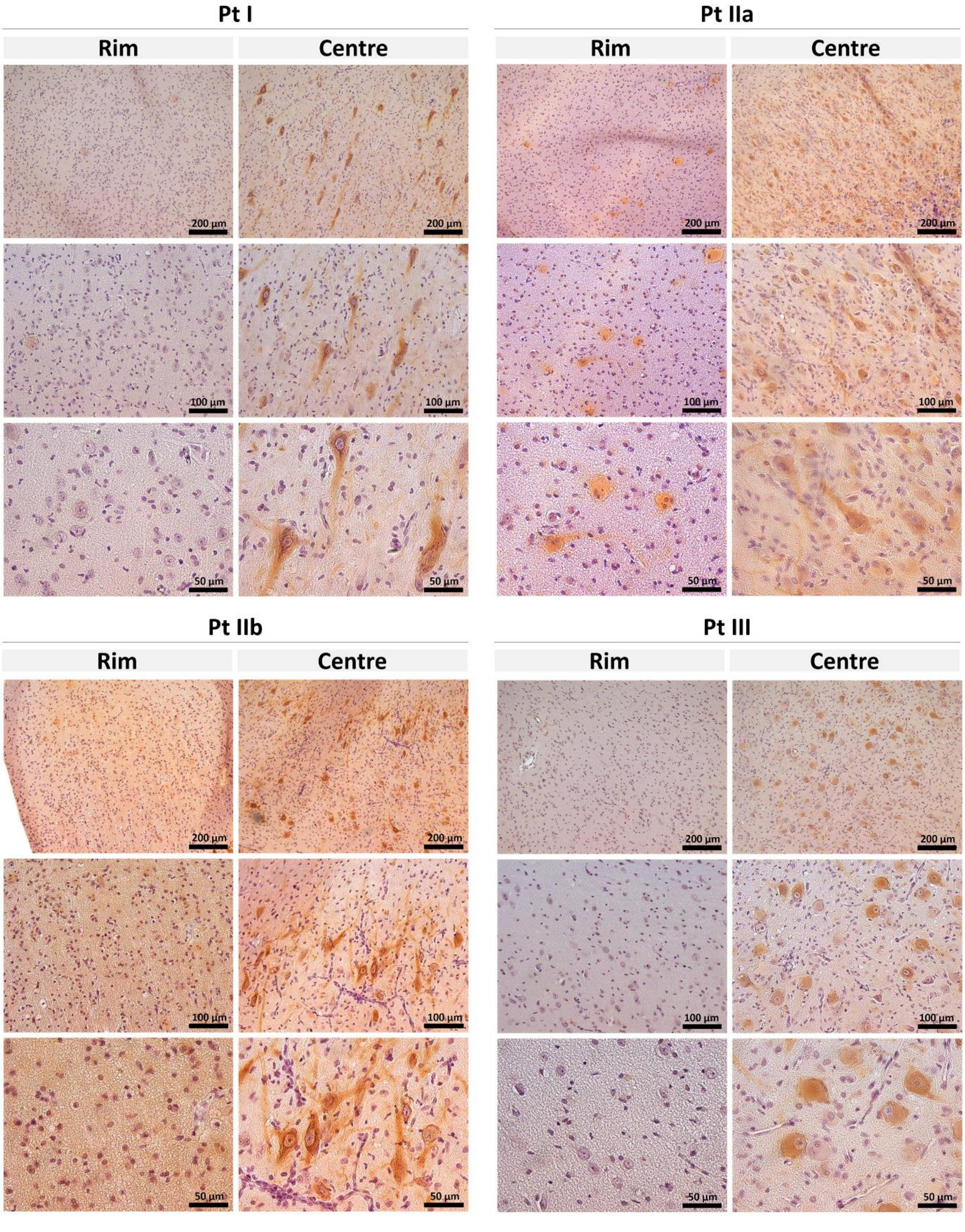


**Supplementary Figure 2: Representative images of Neurofilament IHC of dysmorphic neurons in all patients.** Anti-neurofilament immunohistochemistry staining the dysmorphic neurons in tuber rim and tuber centre demonstrating a typically observed cluster of dysmorphic neurons in the centre biopsy that were absent from the rim. Scale bars are indicated
